# Reporting correct p-values in VEGAS analyses

**DOI:** 10.1101/101014

**Authors:** Julian Hecker, Anna Maaser, Dmitry Prokopenko, Heide Loehlein Fier, Christoph Lange

## Abstract

VEGAS (versatile gene-based association study) is a popular methodological framework to perform gene-based tests based on summary statistics from single-variant analyses. The approach incorporates linkage disequilibrium information from reference panels to account for the correlation of test statistics. The gene-based test can utilize three different types of tests. In 2015, the improved framework VEGAS2, using more detailed reference panels, was published. Both versions provide user-friendly web- and offline-based tools for the analysis. However, the implementation of the popular top-percentage test is erroneous in both versions. The p-values provided by VEGAS2 are deflated/anti-conservative. Based on real data examples, we demonstrate that this can increase substantially the rate of false positive findings and can lead to inconsistencies between different test options. We also provide code that allows the user of VEGAS to compute correct p-values.

---

In 2010, Liu et al. published a gene-based test for genome-wide association studies called VEGAS (versatile gene-based association study)^1^. The approach provides a versatile framework to test genes/genomic regions for genetic association, only requiring summary statistics from single-variant analysis. The approach does not require genotype data. It accounts for linkage disequilibrium (LD) by using simulations from a multivariate normal distribution in which the correlation matrix is defined by the local LD structure. The test is implemented in the software package “VEGAS” and the LD information is extracted from the HapMap reference panel^2^. Recently, Mishra and MacGregor^3^ extended and improved “VEGAS” by incorporating the 1,000 Genomes data as the reference panel^4^ for the local LD structure (“VEGAS2”).

The new software package “VEGAS2” builds on the VEGAS-methodology. Both tools provide the option that only a specific percentage of the most promising SNPs, i.e. the SNPs with the strongest association test signals, are included in the gene-based test. In this case, the observed gene-based statistic in VEGAS is the sum of the squared single-variant association test for the most promising SNPs. To calculate the empirical/permutation-based p-value of this test statistic, in each replicate of the simulations, the sum of the top percentage of the simulated, squared z-scores is computed and compared to the observed statistic.

However, in the software implementation of VEGAS as well as VEGAS2, the top percentage of the signed z-scores is determined and the sum is taken over the squared values. The corresponding R function in the code of VEGAS is

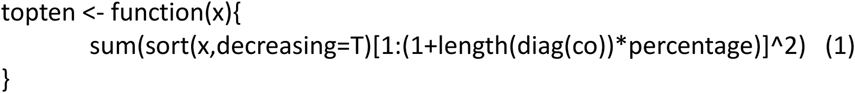

In order to compute the correct test statistic in each replicate, the function should be defined as

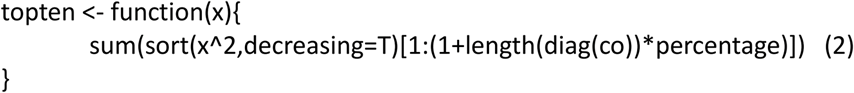

The current implementation (1) therefore provides incorrect p-values for the test statistic, as replicates that consists of large Z-scores with different signs contribute only their largest, positive Z-scores to the computation of the empirical p-value. The correct implementation (2) squares the z-scores and then determines the top percentage of the squared statistics. The empirical p-values provided by “VEGAS” are therefore often smaller/anti-conservative than the correct, permutation-based p-values. The implementation of VEGAS2 also uses definition (1) instead of (2).

Since the decision to perform the top percentage test is very reasonable and it is frequently used in substantive papers, we strongly believe that a correction of this software bug is very valuable for applied research. This is especially true, given the high number of citation for VEGAS/VEGAS2 in data analysis papers. On January 27^th^ 2016, we informed Dr. Stuart MacGregor about this mistake by email. Dr. MacGregor replied that it was a bug indeed and assured us that his group would fix this mistake by the next day. However, to the best of our knowledge, there have not been any software update of the VEGAS/VEGAS 2 packages, since we sent the email to Dr. MacGregor which was about one year ago. We think that this is unfortunate, since, as we outline below, our VEGAS2-results for the Bipolar Disorder Meta-Analysis of the PGC clearly show that the current implementation (1) of the VEGAS test statistic can lead to false positive findings, as VEGAS provides p-values that are substantially anti-conservative. In all our analyses, we used the most recent versions of VEGAS/VEGAS2 (12/15/2016).

To illustrate the impact of this software error, we performed a genome-wide gene-based analysis using the results of the Bipolar Disorder Meta-Analysis of the PGC^5^. We performed the analysis with the offline version of “VEGAS2”, as this is the latest version. We used the whole gene-list provided by “VEGAS2”, a flanking sequence of +/- 10kb and considered the top 10% SNP test. For all other parameters the default option was used.

We performed the tests with the original offline version of “VEGAS2” software and a modified version of “VEGAS2” in which we corrected the software bug as described above. The analysis resulted in gene-based p-values for 23,158 genes.

With the exception of 24 genes, the p-value of the original “VEGAS2” implementation (1) was smaller than the correct p-value based on implementation (2). The average deflation factor between both implementations for the p-value computations was ~1.80. For 218 genes, the factor was greater than 3 and the greatest observed factor was 69.93. Clearly, the factor is expected to be greater, if the number of SNPs mapped to the specific gene increases. For “CSMD1”, the gene with the highest factor of 69.93, the number of mapped SNPs within the flanking sequence was 5,559.

For an overall significance level of 5%, we assessed the number of significantly associated genes after Bonferroni correction for n=23,158 genes. While 15 genes achieved genome-wide significance with incorrect implementation (1), only 9 genes were genome-wide significant with the modified/correct implementation (2).

To emphasize the impact of this software bug on the validity of the test results, we analyzed the example input file which is available for download on the “VEGAS” homepage with the web-based version of VEGAS and the latest offline version of “VEGAS”. We computed the top 9%-, top 20%- and the bestsnp-test with the hapmapCEU information. Using the example input file, “VEGAS” maps 4 variants to the RABL4 and 11 variants to the FOXRED2 gene. Given the number of variants in each gene, the top20%-procedure should correspond to the bestsnp-test for RABL4 and the top 9%-test to the bestsnp-test for FOXRED2. In Table 1, we provide the empirical p-values for our analyses. The results of the modified version (using (2) instead of (1)) are in the last column. All results are based on 10^5 simulations, suggesting sufficient number of replicates in order to obtain stable estimate for the p-values.

**Table 1.**
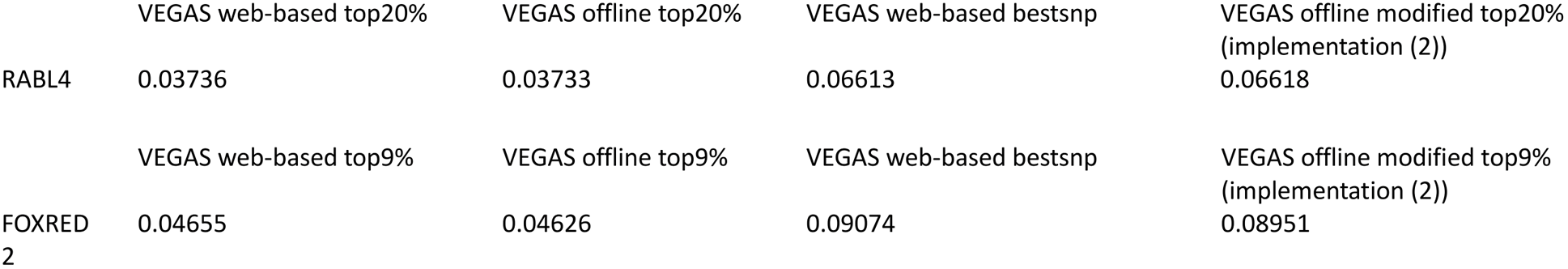

The results of “VEGAS web-based topX%” and “VEGAS offline top X%” match very well, suggesting the same implementation in the web-based verion and in the offline version. However, the results are substantially different from the results obtained by “VEGAS web-based bestsnp”, although theoretically they should be identical. The “VEGAS web-based bestsnp” results agree well with the results of “VEGAS offline modified top X%”. In conclusion, the results of Table 1 proof that the p-values provided by VEGAS are too small, and that the software bug still exists in the web-based and offline version of “VEGAS”.

In this communication, we described why the software implementation of the popular gene-based test frameworks VEGAS and VEGAS2 for the top percentage option is incorrect. Since the top percentage test is frequently applied, we examined the impact of the incorrect computation of the empirical p-value in the top 10% gene-based test statistic by application to the PGC meta-analysis results for bipolar disorder. As the p-values were substantially deflated by the incorrect implementation, VEGAS2 provided false-positive results for 6 genes. Furthermore, we demonstrated inconsistencies between the bestsnp- and the top-percentage-test that also arise from the incorrect implementation. Given the consequences/implication of false-positive results, we strongly believe that user of VEGAS/VEGAS2 should replace (1) in their software copy by the implementation that we provided in (2) so that in the literature correct p-values can be reported.

